# Mitochondrial DNA and temperature tolerance in lager yeasts

**DOI:** 10.1101/391946

**Authors:** EmilyClare P. Baker, David Peris, Ryan V. Moriarty, Xueying C. Li, Justin C. Fay, Chris Todd Hittinger

**Affiliations:** Laboratory of Genetics, Genome Center of Wisconsin, Wisconsin Energy Institute, J. F. Crow Institute for the Study of Evolution, University of Wisconsin-Madison, Madison, WI, USA.; Microbiology Training Program, University of Wisconsin-Madison, Madison, WI, USA.; DOE Great Lakes Bioenergy Research Center, University of Wisconsin-Madison, Madison, WI, USA.; Department of Food Biotechnology, Institute of Agrochemistry and Food Technology (IATA), CSIC, Paterna, Valencia, Spain.; Molecular Genetics and Genomics Program, Washington University, St. Louis, MO, USA.; Department of Genetics, Washington University, St. Louis, MO, USA.; Center for Genome Sciences and System Biology, Washington University, St. Louis, MO, USA.; Department of Biology, University of Rochester, Rochester, NY, USA.

**Keywords:** *Saccharomyces*, evolutionary genetics, mitochondria, thermotolerance, cryotolerance, lager-brewing

## Abstract

A growing body of research suggests that the mitochondrial genome (mtDNA) is important for temperature adaptation. In the yeast genus *Saccharomyces*, species have diverged in temperature tolerance, driving their use in high or low temperature fermentations. Here we experimentally test the role of mtDNA in temperature tolerance in synthetic and industrial hybrids (*Saccharomyces cerevisiae* x *Saccharomyces eubayanus*, or *Saccharomyces pastorianus*), which cold-brew lager beer. We find that the relative temperature tolerances of hybrids correspond to the parent donating mtDNA, allowing us to modulate lager strain temperature preferences. The strong influence of mitotype on the temperature tolerance of otherwise identical hybrid strains provides support for the mitochondrial climactic adaptation hypothesis in yeasts and demonstrates how mitotype has influenced the world's most commonly fermented beverage.

**One Sentence Summary:** Mitochondrial genome origin affects the temperature tolerance of synthetic and industrial lager-brewing yeast hybrids.

## Main Text

Temperature tolerance is a critical component of how species adapt to their environment. The *mitochondrial climatic adaptation* hypothesis (*1*) posits that functional variation between mitochondrial DNA (mtDNA) sequences (mitotypes) plays an important role in shaping the genetic adaptation of populations to the temperatures of their environments. Clines of mitotypes along temperature gradients and associations between mitotype and climate have been observed for numerous metazoan species, including humans (*1*, *2*). Experiments in invertebrates have demonstrated directly that different mitotypes can alter temperature tolerance (*3*, *4*), and mitotype has been associated with adaption to temperature in natural environments (*1*, *5*).

Recent work has suggested that mitotype can also play a role in temperature tolerance in the model budding yeast genus *Saccharomyces* (*6*–*8*). The eight known *Saccharomyces* species are broadly divided between cryotolerant and thermotolerant species (*9*–*11*). Thermotolerant strains (maximum growth temperature ≥36˚C) form a clade that includes the model organism *Saccharomyces cerevisiae* (*12*), while the rest of the genus is more cryotolerant. Most prior research has focused on thermotolerance or the function of mitochondria under heat stress (~37˚C), on mitotype differences within *S. cerevisiae* (*6*, *8*), or on interspecies differences between *S. cerevisiae* and its thermotolerant sister species, *Saccharomyces paradoxus* (*13*). The genetic basis of cryotolerance in *Saccharomyces* has been difficult to determine using conventional crosses focused on the nuclear genome (*14*–*16*). Given how common mitochondrial adaption to cold conditions is among arctic metazoan species (*17*–*19*), mitotype could also conceivably influence cryotolerance in *Saccharomyces*.

In a companion study, Li et al. found that the parent providing mtDNA in hybrids of *S. cerevisiae* and the cryotolerant species *Saccharomyces uvarum* had a large effect on temperature tolerance (*20*). Since *Saccharomyces eubayanus* is the sister species of *S. uvarum* but ~7% genetically divergent, we wondered whether the effect of mitotype would extend to industrial hybrids of *S. cerevisiae* x *S. eubayanus*, sometimes called *Saccharomyces pastorianus* (*21*). While *S. cerevisiae* is well known for its role in human-associated fermentations, it is generally not used to produce lager-style beers, which are brewed at colder temperatures than *S. cerevisiae* can tolerate. Instead, the world's most commonly fermented beverage is brewed using cryotolerant *S. cerevisiae* x *S. eubayanus* hybrids (*21*) that inherited their mtDNA from *S. eubayanus* (*22*, *23*). The recent discovery of *S. eubayanus* (*21*) has sparked substantial interest in understanding the genetics of brewing-related traits to understand how lager strains were domesticated historically and to develop novel lager-brewing strains (*24*–*28*).

To establish the temperature tolerance of *S. cerevisiae* and *S. eubayanus,* relative growth scores were calculated at temperatures ranging from 4-37˚C. Two strains of *S. cerevisiae* (a laboratory strain and a strain used to brew ale-style beers) and two strains of *S. eubayanus* (a derivative of the type strain from Patagonia (*21*) and a strain isolated from North Carolina that is closely related to the ancestor of lager yeasts (*29*)) were tested. Strains were spotted onto plates containing either glucose, a fermentable carbon source, or glycerol, a non-fermentable carbon source that requires respiration to assimilate.

*S. eubayanus* and *S. cerevisiae* had reciprocal temperature responses. *S. eubayanus* strains grew at all temperatures, except 37˚C, while *S. cerevisiae* strains began to decline in relative growth at 15˚C and were completely unable to grow at 4˚C (Fig. 1A-B, Fig. S1-4). Strain-specific differences were also apparent. The *S. cerevisiae*-laboratory strain (*Sc*) and the *S. eubayanus*-North Carolinian strain (*SeNC*) grew relatively weakly compared to conspecific strains. For *Sc,* poor growth was likely driven by auxotrophy, but the reason for *SeNC’s* poor performance is unknown.

**Fig. 1.**
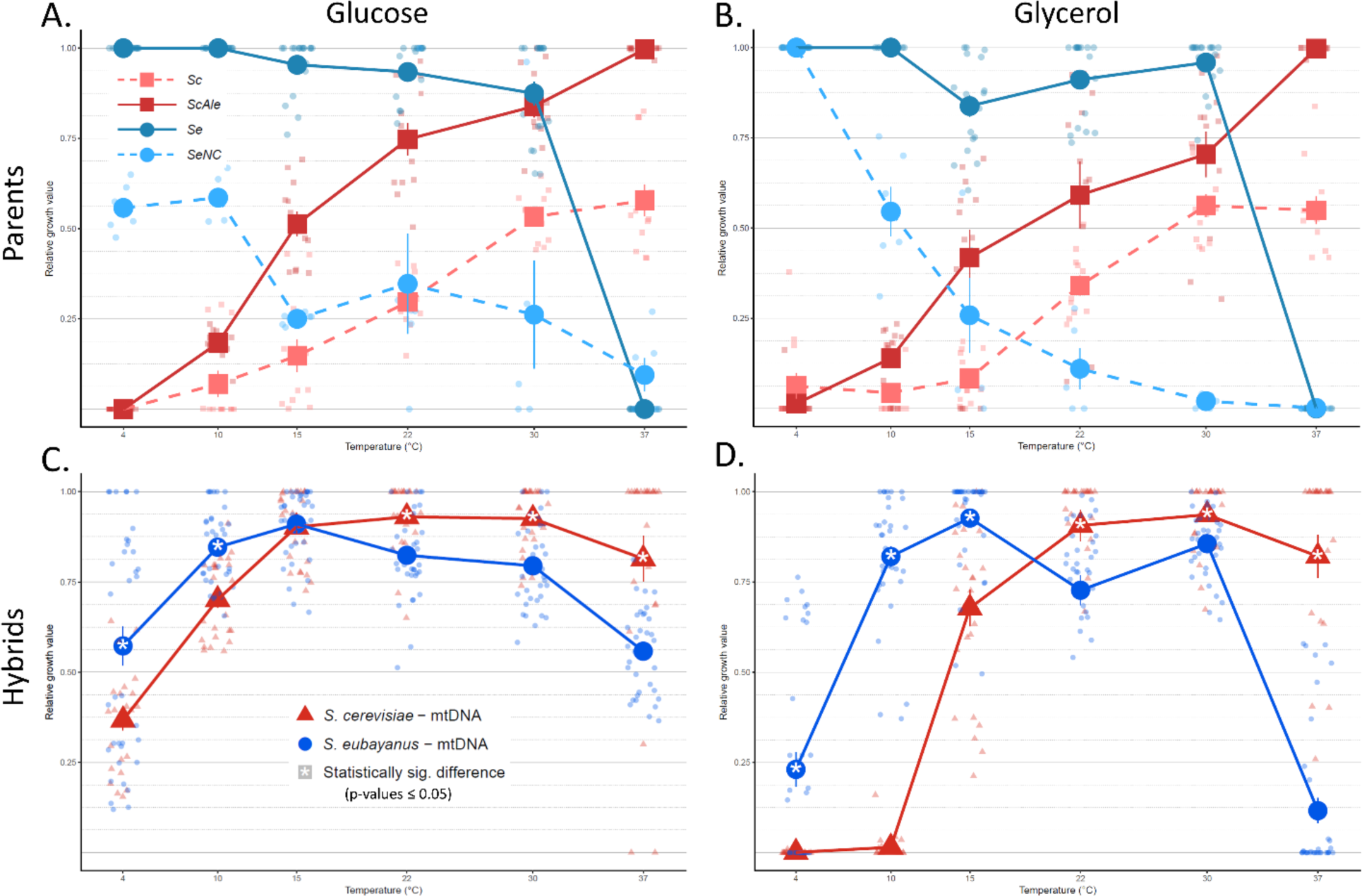
Mitotype affects temperature tolerance in synthetic lager hybrids. Relative growth scores of strains from 4-37˚C combined from all tests. A) On glucose and B) glycerol, relative growth of parent strains carrying their native mtDNA. Parent strains are: *S. cerevisiae*-laboratory strain (*Sc*), *S. cerevisiae*-ale strain (*ScAle*), *S. eubayanus*-type strain (*Se*), and *S. eubayanus*- North Carolinian strain (*SeNC*). C) On glucose and D) glycerol, relative growth of *S. cerevisiae* x *S. eubayanus* synthetic hybrids carrying the mtDNA of different parents, engineered as in Fig. S5. Error bars represent standard error. Differences in relative growth between hybrids carrying different parental mtDNA with p-values of <0.05 were considered statistically significant and are represented by an asterisk. Parents were not tested for significant differences.

To directly test the role of mtDNA, we constructed a panel of synthetic hybrids of *S. cerevisiae* x *S. eubayanus*, controlling the source of mtDNA (Fig. S5). Hybrids tolerated an increased range of temperatures compared to their parents, regardless of mitotype (Fig. 1C-D, Fig. S1-4). These results support a strong role for the nuclear genome in temperature tolerance and indicate some level of codominance between alleles supporting thermotolerance and cryotolerance.

Despite robust growth across temperatures, synthetic hybrids with different mitotypes displayed clear and consistent differences in relative growth. At higher temperatures, *S. cerevisiae* mitotypes permitted increased growth relative to *S. eubayanus* mitotypes, while the same was true for *S. eubayanus* mitotypes at lower temperatures. Relative growth was typically high for both mitotypes on glucose, but significant differences were detected at 5 of 6 temperatures (Fig. 1C). On glycerol, the impact of mitotype was exaggerated (Fig. 1D), and the differences in growth were significant at all temperatures. Subtle background-specific effects were also observed, including a growth defect at 37˚C for the *ScAle* x *SeNC* hybrid carrying *ScAle* mtDNA (Fig. S1).

To test if mtDNA still plays a role in temperature tolerance in industrial lager-brewing hybrids that have been evolving to lagering conditions for many generations, we replaced the native lager mtDNA of *S. eubayanus* origin (*23*) with *S. cerevisiae* mtDNA from *Sc* and *ScAle*, creating lager cybrids (Fig. 2A). Consistent with results for synthetic hybrids, lager cybrids carrying *S. cerevisiae* mtDNA had greater growth at higher temperatures and decreased growth at colder temperatures, especially on glycerol (Fig. 2B-C, Fig. S6). On glucose, strain-specific differences between lager cybrids were particularly apparent. At 30˚C and below, lager cybrids carrying *ScAle* mtDNA grew significantly less than the parental lager strain with its native (*S. eubayanus*) mtDNA (Fig. 2B, Fig. S6A, B), while there was no difference in growth between the parental lager strain and cybrids carrying *Sc* mtDNA, except at temperature extremes (4˚C and 33.5˚C) (Fig. 2B, Fig. S6A, B). On glycerol, both lager cybrids grew significantly less than the industrial strain at 15˚C and below, while they grew significantly more at 22˚C and 30˚C (Fig. 2C, Fig. S6A, C), displaying a shift from lager-brewing toward ale-brewing temperatures. These results show that the strong effect of mtDNA on temperature tolerance seen in synthetic hybrids extends to industrial lager strains under at least some conditions.

**Fig. 2.**
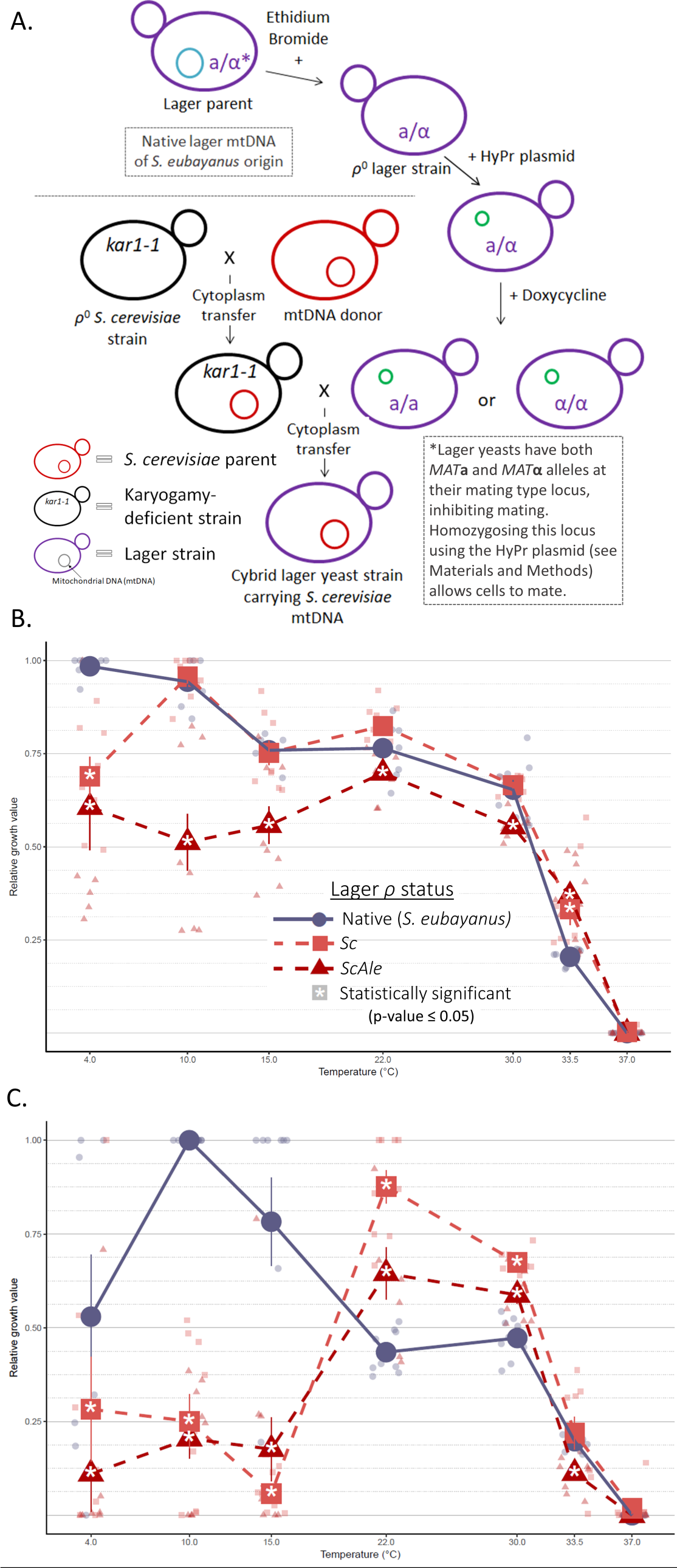
*S. cerevisiae* mtDNA increases the thermotolerance and decreases the cryotolerance of an industrial lager strain. A) Outline of crosses and strain engineering to produce lager cybrids. Yeast cells represent the nuclear genome, large inner circles represent mtDNA, and small green inner circles represent the HyPr plasmid (*30*). Lower case “**a**” and “**α**” indicate mating types. Karyogamy-deficient (*kar1-1*) strains can be of either mating type and are mated to the opposite mating type. Black indicates genetic material from the *S. cerevisiae* karyogamy-deficient strain; red, genetic material from a *S. cerevisiae* parent; blue, genetic material of *S. eubayanus* origin; and purple, a hybrid (i.e. lager) nuclear genome. B) On glucose and C) glycerol, growth of a lager strain with native (*S. eubayanus*) mtDNA and lager cybrids with *S. cerevisiae* mtDNA. Error bars represent standard error, and asterisks indicate statistically significant differences in growth between the cybrid and lager with native mtDNA (p-value <0.05).

We have shown that mtDNA has a significant impact on the temperature tolerance of interspecies hybrids of *S. cerevisiae* and *S. eubayanus*. Along with previous research suggesting hybrid lager yeasts acquired most of their aggressive fermentation traits from *S. cerevisiae* (*25*, *27*, *28*), our results suggest they acquired their cold tolerance from *S. eubayanus* in large part by retaining *S. eubayanus* mtDNA. Our results and methods provide a roadmap for constructing designer lager strains where temperature tolerance can be controlled for the first time (*24*–*28*). Shifting the temperature preference of synthetic or industrial lager strains to warmer fermentation temperatures could substantially reduce the cost of lager brewing by reducing production time and infrastructure requirements. The strain-specific differences observed further suggest that the *S. cerevisiae* parent, the *S. eubayanus* parent, and cytonuclear incompatibilities (*31*), should all be considered during strain construction. Along with the companion study of Li et al. (*20*), the identification of a role for mtDNA in temperature tolerance of these yeasts extends support for the *mitochondrial climatic adaptation* hypothesis (*1*) to fungi and suggests that the outsized role of mtDNA in controlling temperature tolerance may be general to eukaryotes.

## Acknowledgments

The authors thank Thomas D. Fox, Diego Libkind, and José Paulo Sampaio for sharing yeast strains used in this study.

## Author contributions

EPB, DP, XCL, JCF, and CTH conceptualized the study; EPB, DP, and CTH designed experiments; EPB and RVM constructed strains; EPB conducted experiments and analyzed data; DP and CTH supervised RVM; and CTH supervised the project. EPB and CTH wrote the manuscript with input and approval from all authors.

## Data and materials availability

All data are included in the manuscript or its Supplementary Materials. All strains and constructs are freely available for non-commercial research under a material transfer agreement.

## Supplementary Materials

Materials and Methods Figures S1-S6

Tables S1-S2

Supplementary References (*31-54*)

